# Longitudinal differential abundance analysis of microbial marker-gene surveys using smoothing splines

**DOI:** 10.1101/099457

**Authors:** Joseph N. Paulson, Hisham Talukder, Héctor Corrada Bravo

**Affiliations:** Department of Biostatistics and Computational Biology, Dana-Farber Cancer Institute, Boston, MA, USA; Department of Biostatistics, Harvard T.H. Chan School of Public Health, Boston, MA, USA; Applied Mathematics & Statistics, and Scientific Computation graduate program, University of Maryland, College Park, Maryland, USA; Center for Bioinformatics and Computational Biology, University of Maryland, College Park, Maryland, USA; Computer Science Department, University of Maryland, College Park, Maryland, USA

## Abstract

**Background:** High-throughput targeted sequencing of the 16S ribosomal RNA marker gene is often used to profile and characterize the taxonomic composition of microbial communities. This type of big high-through sequencing data is rapidly being applied to various infectious diseases like diarrhea. While many studies are limited to single “snapshots” of these communities, there is increasing recognition that longitudinal profiling of these communities are required to understand community dynamics and the complex relationships between dynamics and phenotypes of interest. Statistical methods that determine microbial features that are differentially expressed are required as an initial step to characterizing phenotypic associations with community dynamics in big data and infectious diseases.

**Results:** We present a novel method for longitudinal marker-gene surveys based on smoothing splines that allows discovery and inference of time periods where specific microbial features are differentially abundant. We applied our method to three 16S marker-gene surveys, including, groups of gnotobiotic mice on two diets, patients challenged with ETEC (H10407), and a vaginal microbiome of healthy women. Employing our methodology we recover known bacterial differences and highlight a few extra species providing insight into when specific changes occurred. Additionally, in the cohort challenged with ETEC we recover proposed probiotic bacteria *Bacteroides xylanisolvens, Collinsella aerofaciens*, and *Faecalibacterium prausnitzii* associatons with healthy individuals.

**Conclusions:** The method presented is, to our knowledge, the first flexible method of its kind implemented as a software capable of detecting time periods of differential abundance for microbial features species between two or more sample groups of interest. Our method is available within the *metagenomeSeq* open-source software for analysis of metagenomic package available through the Bioconductor project and is termed metaSplines.

## Overview

The advent of high-throughput DNA sequencing technology allows scientists to comprehensively examine microbial communities in an ecosystem through targeted sequencing of the 16S rRNA marker-gene (Lindsay et al., 2013). While many studies profile static community “snapshots”, microbial communities do not exist in an equilibrium (Handelsman, Tiedje, & …, 2007). To better understand bacterial population dynamics, many studies are expanding to longitudinal sampling and foregoing cross-sectional or single time-point explorations. Recent studies have characterized healthy microbial communities’ temporal dynamics in the gut (David et al., 2014) and skin following birth (Koenig et al., 2011). Studies have also characterized perturbations to the microbiome due to disease, including, diarrhea (Pop et al., 2014), malnutrition (Smith et al., 2013), SHIV (Morris et al., 2016), and bacterial vaginosis (Ravel et al., 2011). Other studies have explored the effects of external stimuli, including, the effect of diet (Turnbaugh et al., 2009) and antibiotic use (Pop et al., 2016a; Theriot et al., 2014).

With a decrease in sequencing costs more longitudinal data will be generated for varying communities of interest. While data generation will present fewer difficulties, there remain several statistical challenges involved in analyzing these longitudinal datasets. The usual approach in the marker-gene survey literature is to perform pairwise differential abundance tests between specific time points and visually confirm, sometimes using smoothing methods like splines to aid display, how differences are manifested across time (Dickson et al., 2014; Kostic et al., 2015; Seto, Jeraldo, Orenstein, Chia, & DiBaise, 2014). These methods require that analysts provide one or more specific time points to test, and the statistical inferences derived from these procedures are specific to these pairwise tests. Other standard methods for longitudinal analysis test for global differences across time, sometimes using non-linear methods including splines to capture dynamic profiles across time (Smyth, 2005). In this case, statistical inferences are about global changes and not about specific time periods or intervals where differential abundance is detected. An approach that is able to perform statistical inferences about differential abundance over apriori unspecified time periods would provide a more specific view of microbial dynamics for longitundal surveys.

Smoothing spline regression models (G Wahba, 1990) are commonly used to model longitudinal data and form the basis for methods used in a large number of applications (Bravo, 2008; Harezlak, Naumova, & Laird, 2007). Specifically, the Smoothing-Spline ANOVA (SS-ANOVA) method (Gu, 2013) is capable of directly estimating species’ abundances as smooth functions while incorporating sample characteristics as covariates in these models, e.g., sex and age in population studies, or technical factors like processing batches in the chosen model. Incorporating confounding sources of variability, both biological and technical is essential in high-throughput studies (Leek et al., 2010) and require statistical methods capable of estimating both smooth functions and sample-specific characteristics.

In this paper we present a method based on SS-ANOVA for the analysis of longitudinal microbial marker-gene surveys. It is based on a number of important features (i) it incorporates a normalization method designed for these types of surveys (Paulson, Stine, Bravo, & Pop, 2013), (ii) uses semi-parametric modeling to allow incorporation of experimental confounders, essential for large observational studies or studies with complex experimental designs, (iii) it allows discovery of time intervals of differential abundance across multiple phenotypes of interest, and (iv) uses a permutation-based approach to provide robust statistical inferences over discovered time intervals of differential abundance. We have included this methodology in our open source *metagenomeSeq* toolkit for metagenomic data analysis, freely available through the Bioconductor project available at http://bioconductor.org/packages/release/bioc/html/metagenomeSeq.html (Paulson, Talukder, Pop, & Bravo, 2014).

We begin with a brief overview of SS-ANOVA and our general framework followed by the analysis of three marker-gene surveys, including, (i) a gnotobiotic mouse study on differing diets (ii), a cohort of patients challenged with enterotoxigenic *Escherichia coli* (ETEC) and subsequent ciprofloxacin treatment, and (iii) healthy women’s vaginal microbiome over multiple weeks. We highlight temporal dynamics that occur in our reanalysis of the gut microbiomes in shifting diets. In the cohort of patients challenged with ETEC we show the utility of our method in recovering expected a growth in *Escherichia coli* and find associations of potentially probiotic bacteria to individuals that do not become infected with diarrhea. In the vaginal microbiome we illustrate how the SS-ANOVA method captures and incorporates significant background periodic trends in abundances.

### Background on Smoothing Spline ANOVA models

Smoothing Spline analysis of variance (SS-ANOVA) (Grace Wahba, Wang, Gu, Klein, & Klein, 1995) is a semi-parametric method that models data generated from a smooth function *f*(*x*) by assuming that *f* is a function in a Reproducible Kernel Hilbert Space. *f* has a semi-parametric form given by 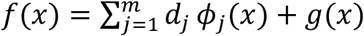 for coefficients *d_j_*, where functions *ϕ_j_* have a parametric form and *g*(*x*) is defined by 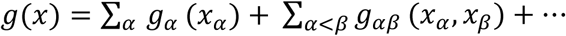 where *g_α_* and *g_αβ_* satisfy the standard ANOVA side conditions. *g_α_* are main effects in the model and *g_αβ_* are the interactions in the model.

The SS-ANOVA estimate of *f*, given data (*x_i_*,*y_i_*), *i* = 1,…, *n*, is a solution of the penalized problem,

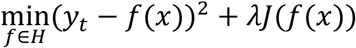

where the first term discourages the lack of fit of *f* and the second term penalizes the complexity of *f* with smoothing parameter *λ* controlling the trade-off between the two. We use Generalized Approximate Cross-Validation (GACV), an approximation to the leave-one-out estimate of the comparative Kullback-Leibler distance between *f̂* and the unknown true *f* to select the regularization parameters used in this process. We also provide Bayesian confidence intervals into our estimation procedure. Further details are provided in the appendix.

### Smoothing Spline Longitudinal Differential Abundance Methodology

In general, we model data in the following form:

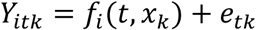

where *i* represents group factor (health status, diet, etc.), *t* represents time, *k* represents replicate observations, *x_k_* are covariates for sample *k* (including an indicator for group membership *I*{*k* ∈ *i*}) and *e_tk_* are independent *N*(0, *σ^2^)* errors. We assume *f_i_* to be a smooth function, defined in an interval [*a, b*], that can be parametric, non-parametric or a mixture of both.

Our goal is to identify time intervals where the absolute difference between two groups 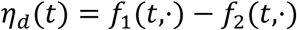 is large, that is, intervals, *R*_*t*_1__,*R*_*t*_2__, where: 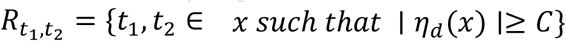 and *C* is a predefined difference threshold.

We applied the SS-ANOVA model to time interval finding by modeling *f* as semiparametric function:

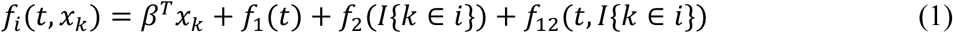

where *β* are coefficients of a linear model of sample covariates (e.g., age, sex), *f*_1_ is the main smooth function over time, *f*_2_ is the main effect term for group *i* and *f*_12_ is a smooth function indicating an interaction term between group membership and time. By encoding group membership using a 0-1 binary variable, the ANOVA side conditions imply that we can directly estimate the difference function 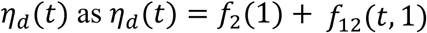. In contrast to other methods, we are able to directly estimate *η_d_*. We use Bayesian confidence intervals above to extend the definition of candidate time intervals of differential abundance *R_t_*_1_,_*t*2_ from before as:

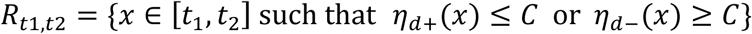

where *η_d+_* and *η_d−_* are the upper and lower 95% confidence intervals. We use this direct estimate of the difference function *η_d_* (*t*) to calculate area statistics 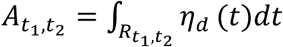 for each time interval of differential abundance. Figure 1 provides an illustrative example of the difference function and test statistic.

**Figure 1.**
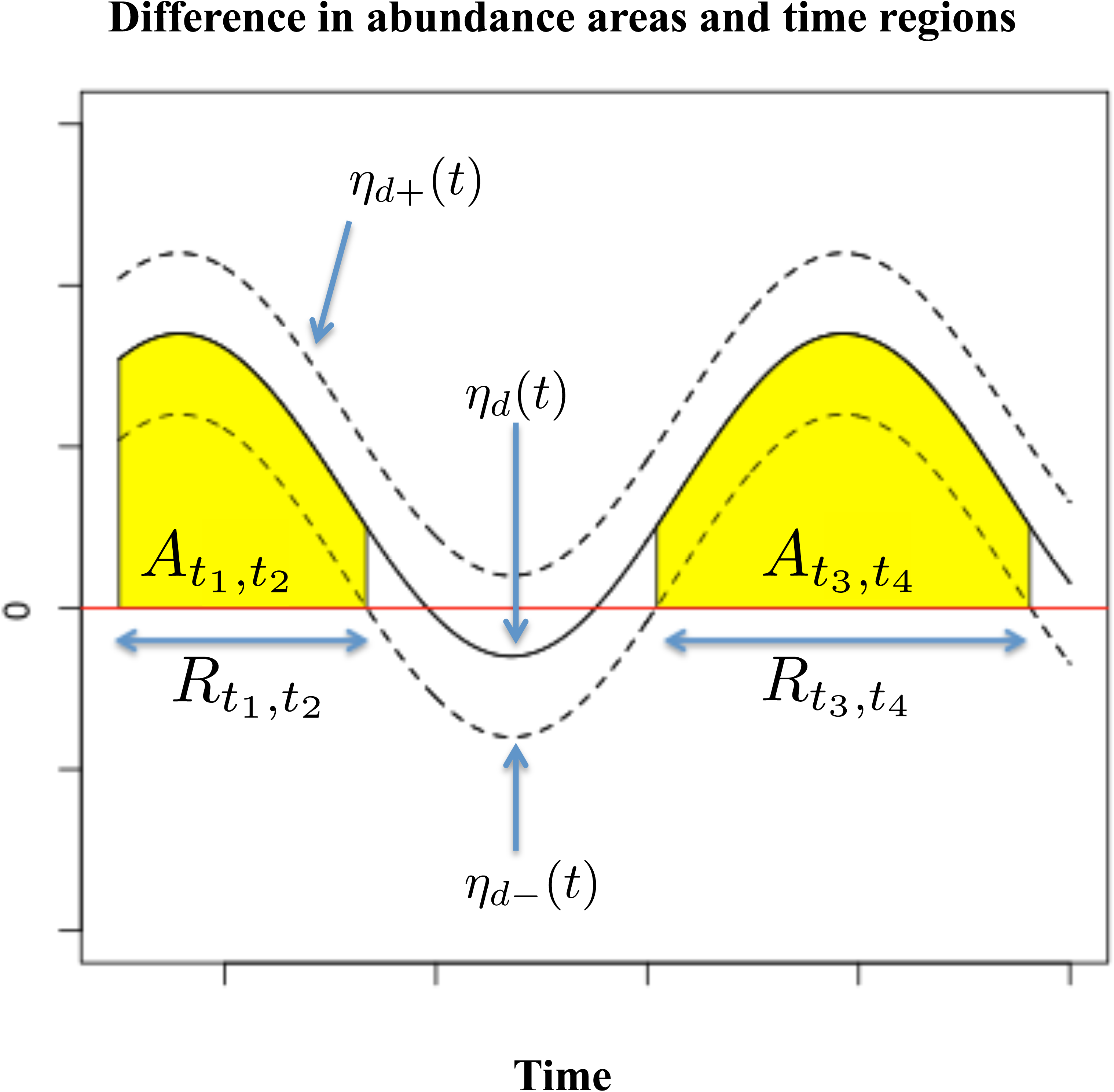
Illustrative example of time intervals of differential abundance. This example shows the difference function, *η_d_* (*t*), with confidence intervals. We choose intervals labeled *R*_*t*_1__,*R*_*t*_2__ and *R*_*t*_3__,*R*_*t*_4__ as possible locations where there are significant difference in response between two groups. The areas under the curve in these intervals, *A*_*t*_1__,*A*_*t*_2__ and *A*_*t*_3__,*A*_*t*_4__, are calculated. These two areas are the test statistic being tested using permutation.

Finally, we contstruct a hypothesis test based on the area statistic to determine time intervals of differential abundance. For this test, the null and alternative hypotheses are:

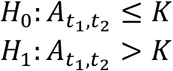

with *K* as a predefined area threshold.

We employ a permutation-based method to calculate a null distribution of the area statistics *A_t_*_1_,_*t*2_'s. To do this, the group-membership indicator variables (0-1 binary variable) are randomly permuted *B* times, e.g., *B =* 1000 and the method above is used to estimate the difference function 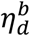 (in this case simulating the null hypothesis) and an area statistics 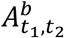 for each random permutation. Estimates 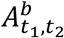 are then used to construct an empirical estimate of *A*_*t*_1__,*A*_*t*_2__ under the null hypothesis. The observed area,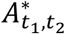, is compared to the empirical null distribution to calculate a *p*-value, i.e.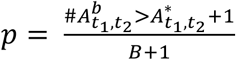. For permutations, we treat negative *η_d_* as negative area and positive *η_d_* as positive area. We adjust for multiple testing across candidate time intervals by using a Bonferroni correction (*α*/n). For example, if we test three candidate time intervals we would reject if the calculated p-value were less than 0.05/3.

## Results

### Smoothing splines analysis of shift in diet increases power

In a study published by Turnbaugh et al., twelve germ-free adult male C57BL/6J mice were fed a low-fat, plant polysaccharide-rich diet. Each mouse was gavaged with healthy adult human fecal material. Following the fecal transplant, mice remained on the low-fat, plant polysaccharide-rich diet for four weeks. A subset of 6 were switched to a high-fat and high-sugar diet for eight weeks. Fecal samples for each mouse went through PCR amplification of the bacterial 16S rRNA gene V2 region weekly. Further details of the experimental protocols and data can be found in (Turnbaugh et al., 2009). We employed the SS-ANOVA modeling approach described above in re-analyzing the data and testing bacterial differences across time for the two differing diets. We aggregated counts to the *class* taxonomic level following CSS normalization.

Using SS-ANOVA we tested the hypothesis that there was no difference in abundance for any particular class due to diet. We considered each bacteria independently of one other. We used SS-ANOVA to estimate abundance of bacteria with the following model:

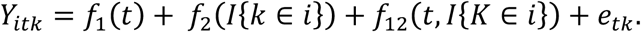

In this application *f*_1_(*t*) represents the effect of time, *f*_2_(*I*{*k* ∈ *i*}) represents the effect of diet and *f*_12_(*t*, *I*{*k* ∈ *i*}) represents the interaction of diet and time. In calculating our test statistic we estimate *η_d_*, a function of the difference in abundance obtained from estimated functions *f*_2_ and *f*_12_ along with a point-wise 95% confidence interval. Using this confidence interval we calculate the difference area for time intervals to detect those above 0.3.

In comparing the two diets a number of bacteria were differentially abundant including *Actinobacteria* prior to multiple-testing correction. We found at least one differential interval in the following classes of bacteria: *Bacilli, Bacteroidetes, Erysipelotrichi*. We observe that in analyzing *Bacteroidetes* with this framework revealed a significant increase immediately after the switch in diet followed by a significant decrease for the duration of the diet. In addition to what was reported by Turnbaugh et al. we were able to uncover greater abundance in Western diets for *Deltaproteobacteria* and *Actinobacteria* for a period immediately following the shift in diet before returning to stability (Figure 2, Table 1).

**Figure 2.**
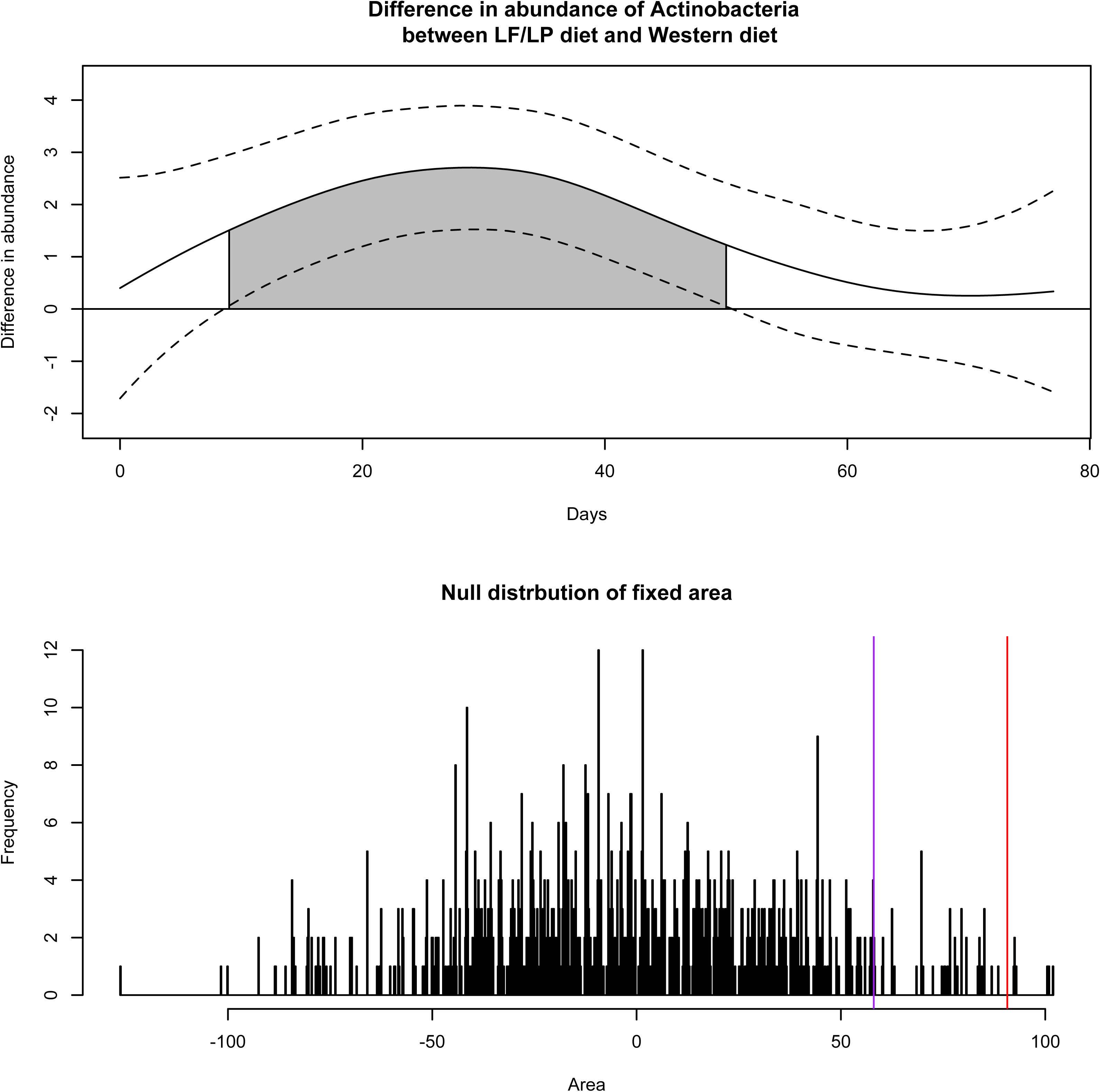
*Actinobacteria* is differentially abundant for a period of time before returning to stability. The top part of the figure is the estimated *η_d_* (*t*) from our model of difference of abundance. Using the Bayesian confidence interval we pick the interval (in grey) where we think there is a significant difference in abundance between the two diets for *Actinobacteria*. The bottom panel reveals the permutation of the null distribution of calculated areas. We show the predicted area in red revealing a significant difference at an alpha level of 0.05.

**Table 1.** Longitudinal differential abundance analysis of multiple diets. Results of metagenomic data using the function metaSplines as stated above. The adjusted P-values are used to reject the null hypothesis. We use *α*=0.05 as a threshold to reject the null hypothesis. A positive area corresponds to a positive shift in abundance for western diet and a negative area corresponds to a positive shift in abundance for LF/LP diet. The diet was switched to western for half the mice after day 21.

| Class names | Interval start | Interval end | Area | p-value | Adjusted p-value |
| --- | --- | --- | --- | --- | --- |
| *Bacteroidetes: interval 1* | 18 | 20 | 6.189 | 0.001 | 0.006 |
| *Bacteroidetes: interval 2* | 22 | 77 | -117.998 | 0.002 | 0.012 |
| *Bacilli* | 21 | 77 | 472.288 | 0.001 | 0.006 |
| *Erysipelotrichi* | 13 | 77 | 118.463 | 0.001 | 0.006 |
| *Deltaproteobacteria* | 20 | 77 | 116.940 | 0.003 | 0.018 |
| *Actinobacteria* | 12 | 47 | 84.132 | 0.005 | 0.030 |

### Smoothing splines ANOVA accurately recalls patient challenge to enterotoxigenic *Eschericia coli*

Diarrhea contributes significantly to the mortality in young children and infants in developing countries (Pop et al., 2014). Approximately 131,000 deaths per year are attributed to enterotoxigenic *Escherichia coli* infection as well as an estimated 10 million cases of travelers’ diarrhea. To further undertand how the intestinal microbiome is altered during infection, Pop et al. subjected 12 volunteers to ETEC (H10407) and subsequent antibiotic treatment (Pop et al., 2016b). They collected samples pre-infection and 104 samples in the nine days following infection. Of the 12 volunteers, 5 subjects developed severe diarrhea with 7 remaining asymptomatic. We employed the same SS-ANOVA modeling approach described above in re-analyzing the data and testing bacterial differences across time between diseased and healthy patients. We aggregated counts to the *species* taxonomic level following CSS normalization. We considered each species independently.

We chose the following model to test our approach:

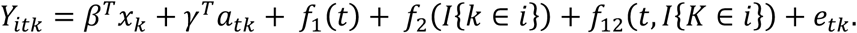

In this application *x_k_* is an indicator for the individual patient as a fixed effect, *a_tk_* is an indicator for the particular samples provided ciprofloxacin treatment, *f*_1_(*t*) represents the effect of time, *f*_2_(*I*{*k* ∈ *i*}) represents the effect for subjects that developed diarrhea after challenge (disease group) and *f*_12_(*t*, *I*{*k* ∈ *i*}) represents the interaction of disease group and time. As before, we estimate *η_d_*, a function of the difference in abundance calculating a 95% confidence interval to find difference area for regions above our predefined cutoff of 0.3.

Patients’ gut microbiota was collected from day -1 to 9 with infection at day 0. Only 22/147 species had time intervals of potential differential abundance as estimated with SS-ANOVA. We recovered the expected largest difference in abundance due to a bloom of *Escherichia coli* starting from the day after infection (Figure 3 and Table 2). While a few bacteria were differentially abundant prior to infection (6/17), the majority of bacteria began to reveal a shift in abundance post. Abundant species that were associated with the diseased group included commensal bacteria, *Roseburia Faecis, Roseburia inulinivorans, Bacteroides ovatus*, and *Bacteroides thetaiotaomicron.* These bacteria potentially interact with ETEC or a bi-product of *E. coli* or are less sensitive to ciprofloxacin treatment which occurred earlier for certain diseased patients (Wexler, 2007). Abundant species that were associated with the healthy individuals included, *Alistipes sp., Bacteroides xylanisolvens*, *Collinsella aerofaciens*, and *Faecalibacterium prausnitzii*. *Bacteroides xylanisolvens, Collinsella aerofaciens*, and *Faecalibacterium prausnitzii* have been proposed as probiotics, potentially playing a role in reducing inflammation and acting as a probiotic (Malinen et al., 2010; Miquel et al., 2013; Ulsemer et al., 2012).

**Figure 3.**
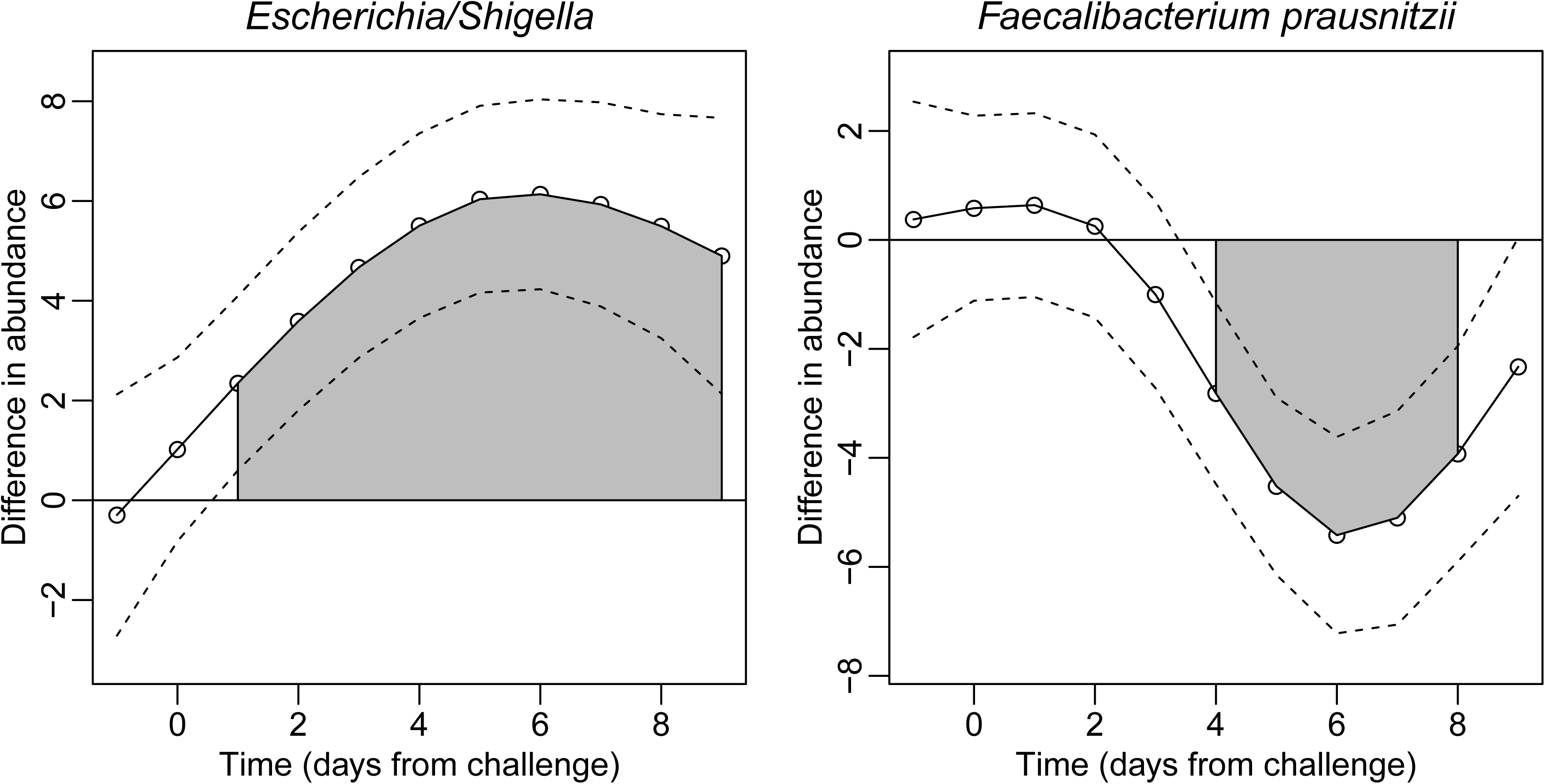
Smoothing Spline ANOVA recovers expected ETEC challenge as well as probiotic bacteria associated with healthy individuals. Estimated *η_d_* (*t*) from our model of difference of abundance for *Escherichia/Shigella* (left) and *F. prausnitzii* (right). We observed that the differential abundance of *Escherichia/Shigella* follows immediately post infection and begins to decay after subsequent antibiotic treatment. Additionally, post challenge and treatment it would appear that there is a greater reduction in *F. prausnitzii* post diarrheal occurrence and antibiotics.

**Table 2.** Longitudinal differential abundance analysis of patients challenged with ETEC. Results of metagenomic data using the function metaSplines as stated above. The adjusted P-values are used to reject the null hypothesis. We use *α*=0.05 as a threshold to reject the null hypothesis. A positive area corresponds to a positive shift in abundance for individuals that eventually became symptomatic and a negative area corresponds to a positive shift in abundance for individuals remaining asymptomatic. All individuals were infected with ETEC.

| Species name | Interval start | Interval end | Area | p-value | Adjusted p-value |
| --- | --- | --- | --- | --- | --- |
| *Bacteroides ovatus* | 5 | 9 | 14.303 | 0.017 | 0.374 |
| *Escherichia/Shigella* | 1 | 9 | 43.66 | 0.001 | 0.022 |
| *Faecalibacterium prausnitzii* | 4 | 8 | -18.107 | 0.001 | 0.022 |
| *Ruminococcus faecis* | 5 | 6 | 1.022 | 0.001 | 0.022 |
| *Bacteroides dorei* | -1 | 4 | 19.692 | 0.001 | 0.022 |
| *Eubacterium rectale* | 5 | 6 | -2.74 | 0.001 | 0.022 |
| *Bacteroides xylanisolvens* | 1 | 9 | -20.243 | 0.001 | 0.022 |
| *Succiniclasticum ruminis* | -1 | 6 | -19.336 | 0.001 | 0.022 |
| *Collinsella aerofaciens* | 2 | 8 | -17.71 | 0.001 | 0.022 |
| *Bacteroides thetaiotaomicron* | 5 | 9 | 13.332 | 0.001 | 0.022 |
| *Alistipes sp.* | 5 | 9 | -13.833 | 0.001 | 0.022 |
| *Bacteroides plebeius* | 4 | 7 | 6.86 | 0.001 | 0.022 |
| *Alistipes putredinis* | 7 | 9 | -7.016 | 0.001 | 0.022 |
| *Ruminococcaceae incertae sedis* | 5 | 9 | -5.005 | 0.001 | 0.022 |
| *Roseburia faecis* | 5 | 9 | -18.032 | 0.001 | 0.022 |
| *Sutterella stercoricanis* | -1 | 4 | -13.931 | 0.001 | 0.022 |
| *Oscillibacter* | 6 | 7 | -1.885 | 0.001 | 0.022 |
| *Roseburia inulinivorans* | 2 | 7 | 16.468 | 0.001 | 0.022 |
| *Eubacterium ventriosum* | 2 | 3 | 1.81 | 0.001 | 0.022 |
| *Romboutsia lituseburensis* | -1 | 0 | 2.447 | 0.001 | 0.022 |
| *Olsenella uli* | -1 | 1 | 2.143 | 0.012 | 0.264 |
| *Turicibacter sanguinis* | -1 | 0 | 1.062 | 0.001 | 0.022 |

### Periodic smoothing splines models cyclical differences in the vaginal microbiome

We observed the utility of a periodic smoothing spline approach to model bacterial abundances and differential abundance estimates that fluctuate through time (Figure 4). Nugent scores are an important measurement in diagnosing women’s health, in particular to bacterial vaginosis (Nugent, Krohn, & Hillier, 1991), and is directly related to the presence of large Gram-positive rods (various *Lactobacillus* morphotypes). We tested the hypothesis that there was no difference in abundance for women that tended to have high to intermediate Nugent scores compared to women with low scores. We chose to separate women that had low nugent scores from those with intermediate to high values. We observed a clear separation following PCA analysis between these two groups of individuals following a PCA analysis. In particular, we highlight the use of periodic smoothing splines on the most abundant organism, *Lactobacillus iners*, from a 2010 study of the vaginal microbiome of reproductive-age women (Ravel et al., 2011). Using a more flexible model and less parameterized model we are able to confirm the fluctuation of *Lactobacillus iners* in healthy and typically stable patient communities of low nugent scores.

**Figure 4.**
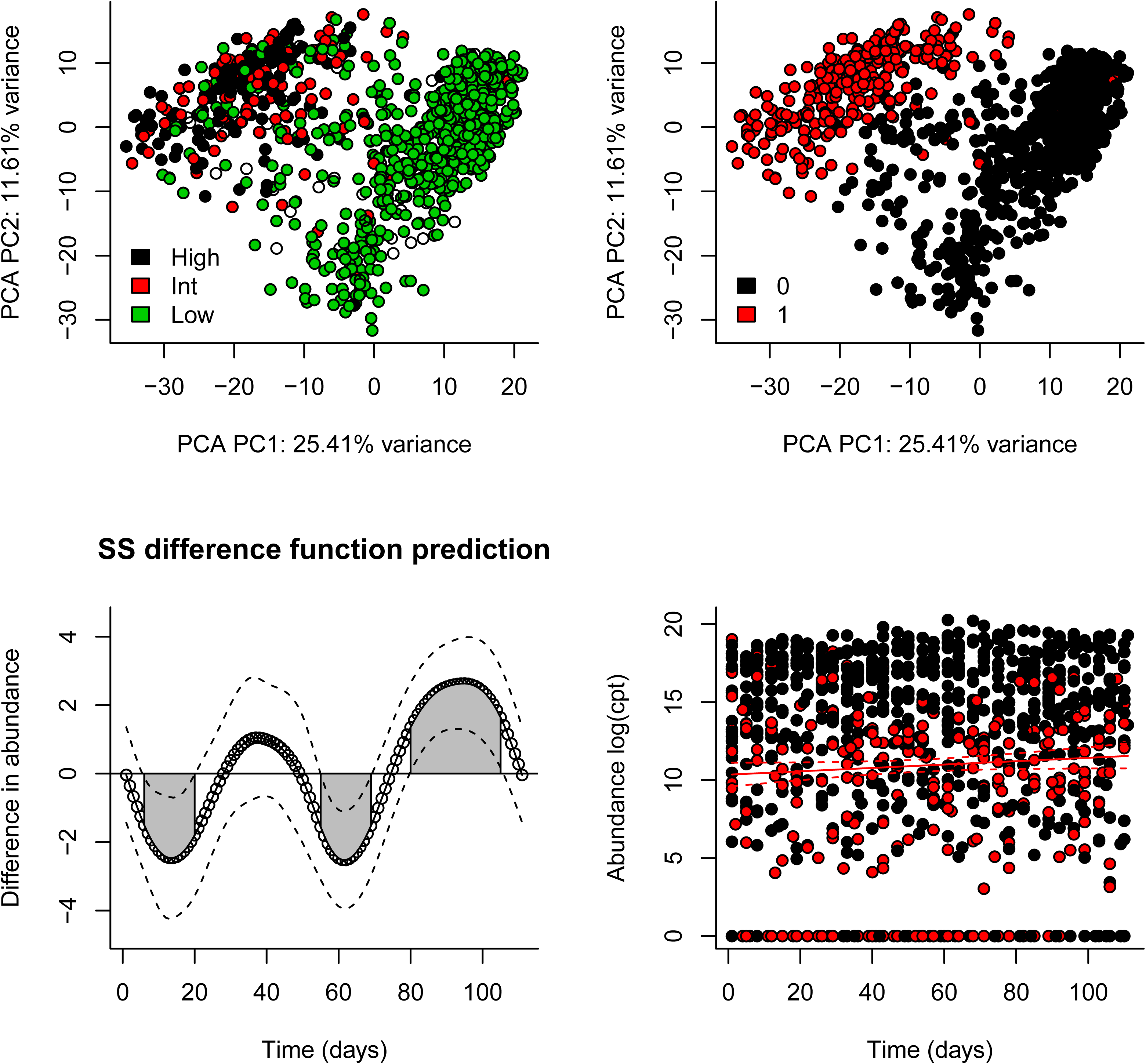
Vaginal microbiome time-series analysis reveals cyclical trend. Top left, PCA analysis of samples from the vaginal microbiome cohorot of Ravel et al. 2010. Colors represent in the top figure nugent scores for samples characterized as either, low, intermediate, or high. Top right, samples are recategorized as 0 - low nugent patients, or 1 high/intermediate nugent patients. Bottom left, Estimated function of difference in abundance for *Lactobacillus iners*. We observed that the differential abundance function follows an approximately monthly cycle. Top right, observed data with average of all data points running through the middle.

### Comparison to alternative methods

We compared the SS-ANOVA permutation based approach to two alternative approaches on the large healthy/malnourished infant cohort and the gnotobiotic mouse diet study. The first method consisted of a pairwise *t*-test for each time point. We did not observe any differential abundance at any time point in the infant cohort due to a lack of power. However, in analyzing the mouse study, *Bacteroidetes, Bacilli, Erysipelotrichi*, was significantly different in both methods between the same time intervals. However, using the SS-ANOVA approach we gained significant intervals of differential abundance for *Deltaproteobacteria* and *Actinobacteria*.

The second alternative method in longitudinal differential abundance analysis is to fit a natural spline and calculate an F-statistic on the two fits. In our analysis of the diet study, the natural spline approach confirmed our analysis, but additionally reported *Epsilonproteobacteria*. This organism was present in only 3 samples at low abundance compared to an average of 119 samples for the other reported bacteria most likely a false positive.

## Methods

### Data acquisition and normalization

Three 16S ribosomal RNA marker-gene surveys were used in the development and analysis of our method, metaSplines. The infant cohort and mouse gut shifting diet cohort datasets were downloaded from Bioconductor with existing annotations that were used to aggregate normalized counts. The ETEC challenged samples are available at: https://bioconductor.org/packages/release/data/experiment/html/etec16s.html. The multiple diet study is available within the *metagenomeSeq* pacakge as an example dataset. The vaginal microbiome count data and annotation was downloaded from the supplementary material of Ravel et al. at http://www.pnas.org/content/suppl/2010/06/03/1002611107.DCSupplemental/st04.xlsx (Ravel et al., 2011). Counts were converted from proportions back to raw counts rounding to the nearest integer. Further details for the sequencing, clustering and annotation are available in (Pop et al., 2016a; Ravel et al., 2011; Turnbaugh et al., 2009).

Data normalization is a crucial initial step in making counts comparable across samples. Counts were normalized per the cumulative sum scaling (CSS) method described in (Paulson et al., 2013). We aggregated normalized counts by annotation to various levels of the taxonomic tree including genera and class levels. Particular levels were chosen for the appropriate comparisons to previously published results. To analyze the infant cohort we aggregated normalized counts to genus level annotations. Classes were analyzed in the diet study and species for the vaginal microbiome cohort.

### Software

The SS-ANOVA based method is available in the Biocondcutor package, *metagenomeSeq*. We provide an extensive documentation and vignette for users to analyze their own datasets. For analyses presented we used *metagenomeSeq* version 1.15.3.

### Conclusions

We proposed a method for the time interval-finding task based on smoothing spline methods that is direct and interpretable. We applied our approach three different microbiome studies, an infant cohort, gnotobiotic mouse longitudinal study and healthy women vaginal consortium. Additionally, we performed a simple comparison analysis on the first two datasets using commonly employed methods, namely a pairwise comparison and F-test on natural spline fits.

The smoothing-spline ANOVA method accurately detects time intervals of differential abundance by directly estimating the difference function of interest. This is the first method specifically developed for testing differentially expressed intervals of marker-gene survey data. As longitudinal data becomes less cost-prohibitive methods to analyze the complex interactions in big infectious microbial data will necessitate methods like the one proposed.

Additionally, Smoothing Spline ANOVA methods are potentially applicable to other high-throughput genomic data. Resolving base-pair differences is an important problem in several other high-throughput genomic data analysis applications including ChIP-seq (number of aligned reads in a given interval), DNA methylation (methylation level at a genomic locus), and recently RNA-seq (number of aligned reads in a genomic position). Intervals of interest in these applications include contiguous genomic intervals in which base-pair level measurements show significant differences between groups of samples. Recent widely used methods for this task take a smoothing approach to find these intervals of significant difference. However, these methods employ an indirect approach that is inefficient and appropriate interpretation of their estimates is not possible. As interval-finding applications continue to flourish with the advent of high-throughput assays, specifically next-generation sequencing, the general methodology presented here will address a rapidly increasing number of critical applications in genomics.

## Acknowledgements

We would like to thank Dr. O. Colin Stine for useful discussions and access to the data performed on the infant gut cohort. This work was funded in part by NIH grants R01 HG006102 and HG005220 to HCB. JNP was funded in part by the US National Science Foundation Graduate Research Fellowship award DGE 0750616 and by NIH R01 HL111759 and NIH U01 CA190234 to John Quackenbush at Dana-Farber Cancer Institute.

